# The pharynx of the iconic stem-group chondrichthyan *Acanthodes* (Agassiz, 1833) revisited with micro computed tomography

**DOI:** 10.1101/2023.08.23.554409

**Authors:** Richard P. Dearden, Anthony Herrel, Alan Pradel

**Affiliations:** Naturalis Biodiversity Centre, Darwinweg 2, 2333 CR, Leiden; CR2P, Centre de Recherche en Paléontologie–Paris, Muséum national d’Histoire naturelle, Sorbonne Université, Centre National de la Recherche Scientifique, Paris cedex 05, France; UMR 7179 (MNHN-CNRS) MECADEV, Département Adaptations du Vivant, Muséum National d’Histoire Naturelle, Paris, France

**Keywords:** acanthodian, branchial skeleton, chondrichthyan, computed tomography, Permian, pharynx

## Abstract

*Acanthodes* has long been the primary source of information on the pharyngeal skeleton of ‘acanthodians’, a stem-group chondrichthyan grade. Because of this its anatomy has played an outsized role in attempts to understand the evolution of the jawed vertebrate pharynx and the clade as a whole. However, the anatomy of the pharynx of *Acanthodes* remains poorly understood and subject to several competing interpretations. We use computed tomography (CT) to image the articulated pharyngeal skeletons of three specimens of *Acanthodes confusus* from Lebach, Germany. *Acanthodes* had a *mélange* of osteichthyan-like and chondrichthyan-like morphologies in its pharyngeal skeleton. Like other stem-chondrichthyans, *Acanthodes* had a basihyal with no hypohyals, and four pairs of posteriorly oriented pharyngobranchials. Like osteichthyans, *Acanthodes* possessed an interhyal, but lacked the separate infra- and supra-pharyngobranchial elements present in osteichthyans and the crown-chondrichthyan *Ozarcus*. Using this new data we build and animated a digital 3D model of the pharyngeal endoskeleton in *Acanthodes*, showing that the jaws would have swung outwards during the opening cycle, increasing the anteriorly facing area of the gape for suspension feeding. These new data provide a more definitive picture of the anatomy of a taxon that has long been of great significance in early vertebrate palaeontology.

## Introduction

The late Palaeozoic stem-group chondrichthyan *Acanthodes* has been a focal point of efforts to understand the evolution of the vertebrate pharynx since the 19^th^ century (Reis, 1890, 1894, 1895, 1896; Dean, 1907; Watson, 1937; Miles, 1973a; Jarvik, 1977; Heidtke, 2011, 2015; Davis *et al*., 2012; Brazeau & de Winter, 2015). The extensive preservation of the delicate and easily disarticulated visceral skeleton in numerous fossils of *Acanthodes confusus* from the Early Permian of Lebach, Germany makes it one of very few keystone Palaeozoic taxa and for a long time the only “acanthodian” stem-group chondrichthyan preserving information on the anatomy of the pharyngeal skeleton. Recently a number of three-dimensionally preserved pharyngeal skeletons of Palaeozoic chondrichthyans have been described in great detail using computed tomographic (CT) methods, including other stem-group chondrichthyans (Coates *et al*., 2018; Dearden *et al*., 2019; Maisey *et al*., 2019) as well as Palaeozoic crown-group members (Pradel *et al*., 2014, 2021; Coates *et al*., 2019, 2021; Frey *et al*., 2019, 2020; Hodnett *et al*., 2021; Klug *et al*., 2023), which show an unexpected range of feeding modes. However, *Acanthodes* remains important as one of the best preserved of these and is further interesting from a functional perspective as a very late-occurring acanthodian, seemingly adapted to be an anguilliform filter feeder, in the broader ecological context of the late Palaeozoic (Stack *et al*., 2020).

Despite there being numerous, well-studied fossils in museum collections preserving the pharyngeal skeleton of *Acanthodes* its anatomy remains uncertain. Interpretations of the pharynx of *Acanthodes* remain based almost entirely (but see (Dearden & Giles, 2021)) on acid-prepared moulds, studied by making casts either physically (Miles, 1973a) or more recently digitally (Brazeau & de Winter, 2015). While many detailed studies of these casts of *A. confusus* have been carried out, the mouldic nature of the specimens have resulted in interpretational ambiguities, further exacerbated by the segmented ossification pattern of the visceral skeleton (Reis, 1896; Dean, 1907; Watson, 1937; Miles, 1964, 1965, 1968, 1973a; Nelson, 1968; Jarvik, 1977)). This has led to conflicting reconstructions of the pharyngeal anatomy of *Acanthodes*, with very different interpretations of characters such as the hyoid arch components and orientation of pharyngeal elements (Fig. 1). Several of the variable features of these reconstructions are potentially important phylogenetic characters and recent reassessment of other aspects of the endoskeletal anatomy of *Acanthodes* have had influential phylogenetic implications (Davis *et al*., 2012; Brazeau & de Winter, 2015). More generally they feed into hypotheses of the origins of gnathostome pharyngeal anatomy (Pradel *et al*., 2014; Coates *et al*., 2018; Dearden *et al*., 2019).

**Figure 1.**
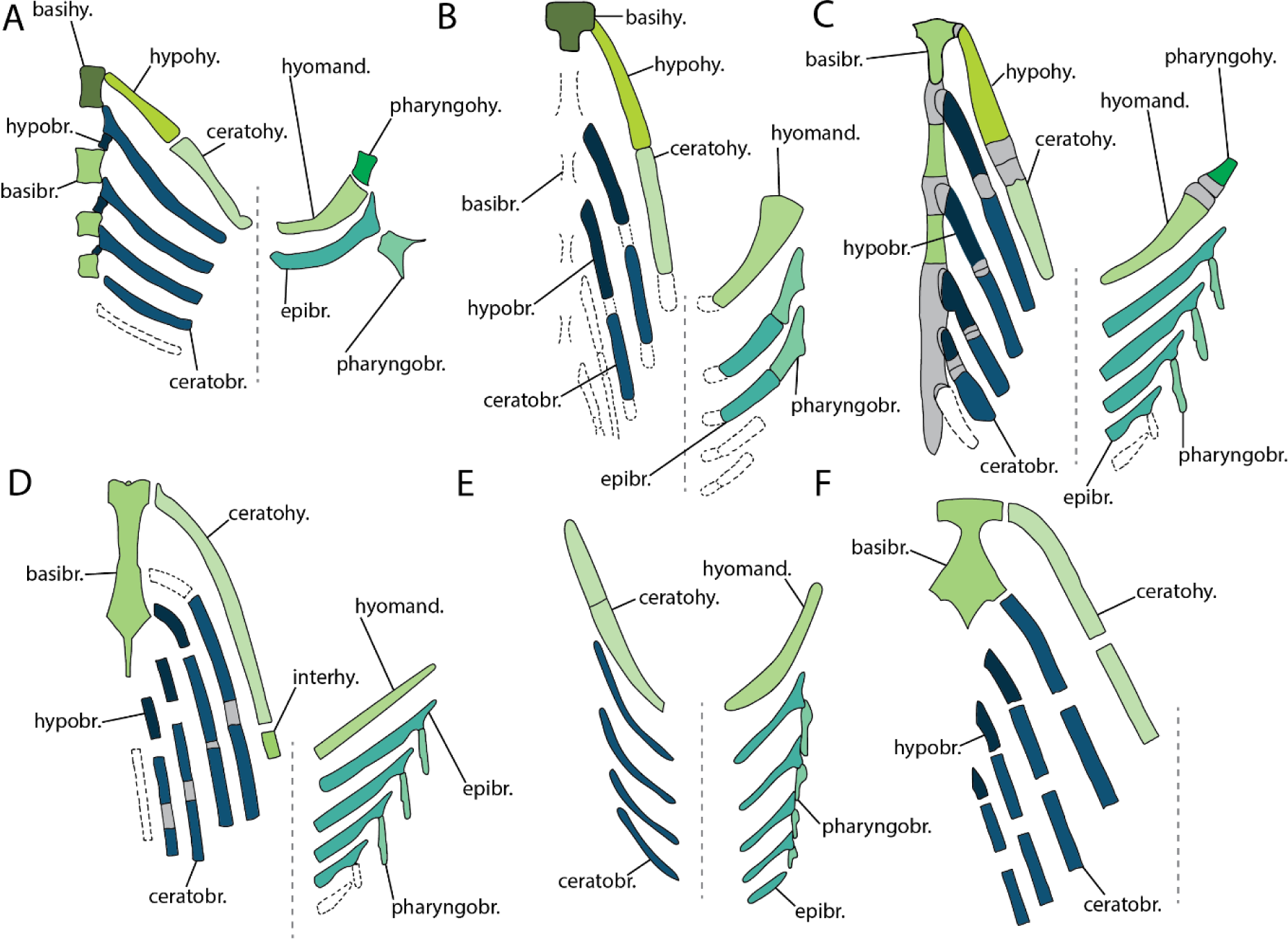
Previous reconstructions of the branchial arch skeleton in *Acanthodes*. A, Reis 1896. B Watson 1937. C, Nelson 1968. D, Miles 1973. E Jarvik 1977. F, Gardiner 1984. Terminology has been standardized to match terms used in this study. Abbreviations: basibr., basibranchial; basihy., basihyal; ceratobr., ceratobranchials; ceratohy., ceratohyal;; epibr,epibranchial; hyomand., hyomandibula; hypohy., hypohyal; interhy., interhyal; pharyngobr., pharyngobranchials; pharyngohy., pharyngohyal. Matching colours denote serially homologous elements. Grey indicate inferred areas of cartilage. Black dashed lines indicate possible elements. Grey dashed line indicates junction of dorsal and ventral branchial skeleton.

Here we aim to resolve uncertainty in the pharyngeal anatomy of *Acanthodes confusus* carrying out a detailed redescription. Rather than using mouldic specimens, we do this by using CT scanning to image concretions from Lebach which have not been acid prepared. Using the resulting 3D models we reconstruct the anatomy of its visceral skeleton, reappraise this anatomy and its functional morphology in the context of what is now known about the pharyngeal anatomy of Palaeozoic gnathostomes.

## Material and Methods

### Specimens

In this study we describe three specimens of *Acanthodes confusus* from the collections of the Muséum National d’Histoire Naturelle (MNHN), Paris: MNHN SAA21 (Fig 2, and MNHN SAA20 (Fig 3), MNHN SAA24 (Fig 4). The majority of Lebach vertebrate remains have been prepared using acid to create moulds, so that they can be studied using peels and casts Miles 1973(Miles, 1973a). We selected these three because they visibly preserve articulated visceral skeletons which have not been extensively acid prepared. In the MNHN collections these specimens are referred to as *Acanthodes bronni*; the well-ossified *Acanthodes* taxon from Lebach has since been revised as and placed in the species *Acanthodes confusus* (Heidtke, 2011; Coates *et al*., 2018).

**Figure 2.**
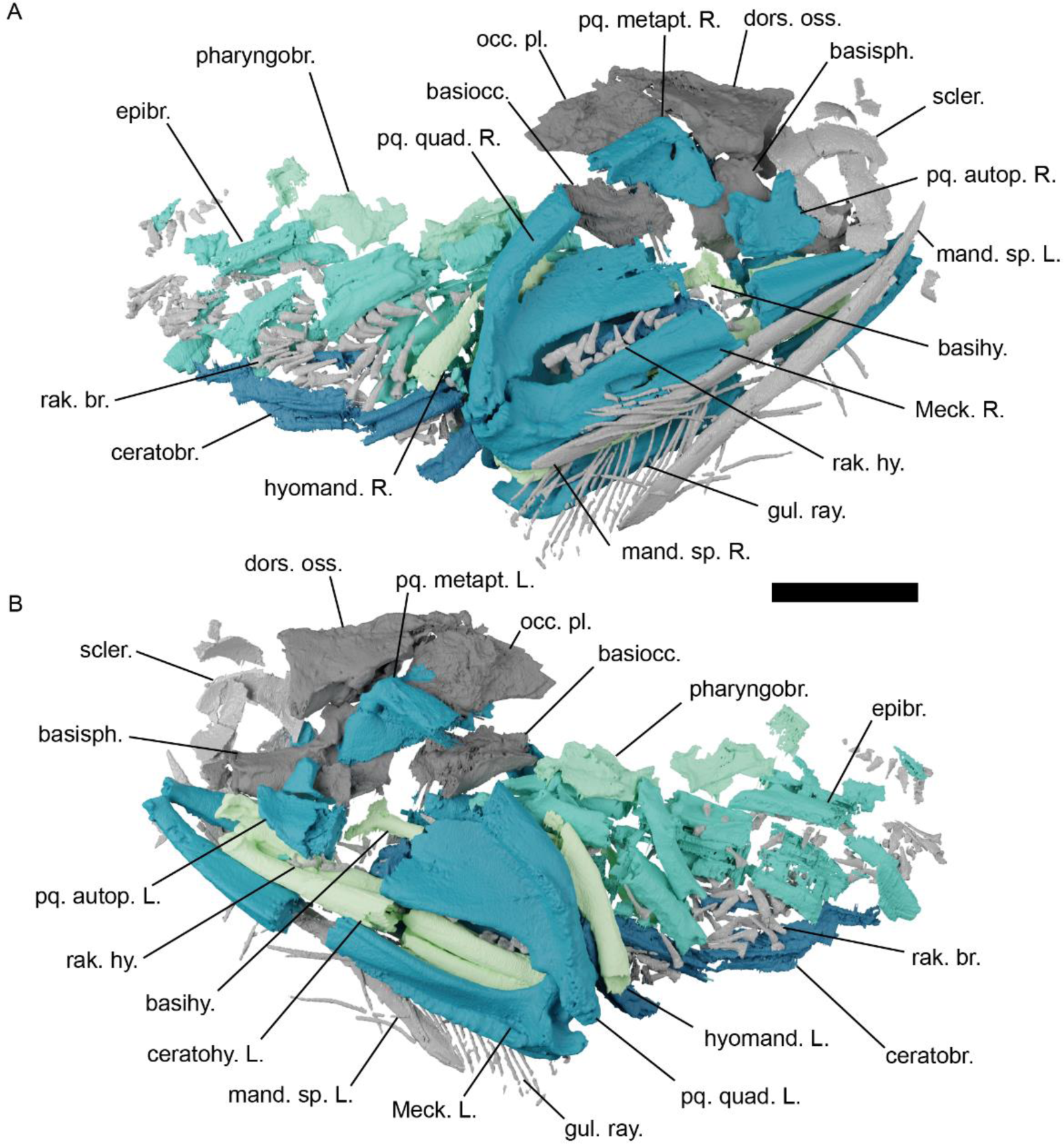
The head skeleton of *Acanthodes confusus* MNHN SAA21, visualised using computed tomography. A, in right lateral view. B, in left lateral view. Abbreviations: basihy., basihyal; basiocc., basioccipital; basisph., basisphenoid; ceratobr., ceratobranchials; ceratohy., ceratohyal; dors. oss., dorsal ossification of neurocranium; epibr,epibranchial; gul. ray., gular rays; hyomand., hyomandibula; L., left; mand. sp., mandibular splint; Meck., Meckel’s cartilage; occ. pl., occipital plate; pharyngobr., pharyngobranchials; pq. autop., palatoquadrate autopalatine; pq. metapt., palatoquadrate metapterygoid; pq. quad., palatoquadrate quadrate; R., right; rak.br., branchial rakers; rak. hy., hyoid rakers; scler, sclerotic ring. Colour scheme: blue-greens, elements of visceral endoskeleton with mandibular arch, hyoid arch, hypobranchials, ceratobranchials, epibranchials, and pharyngobranchials coloured independently; dark grey, cranial and pectoral endoskeleton; light grey, elements of the dermal skeleton. Scale bar = 20 mm.

**Figure 3.**
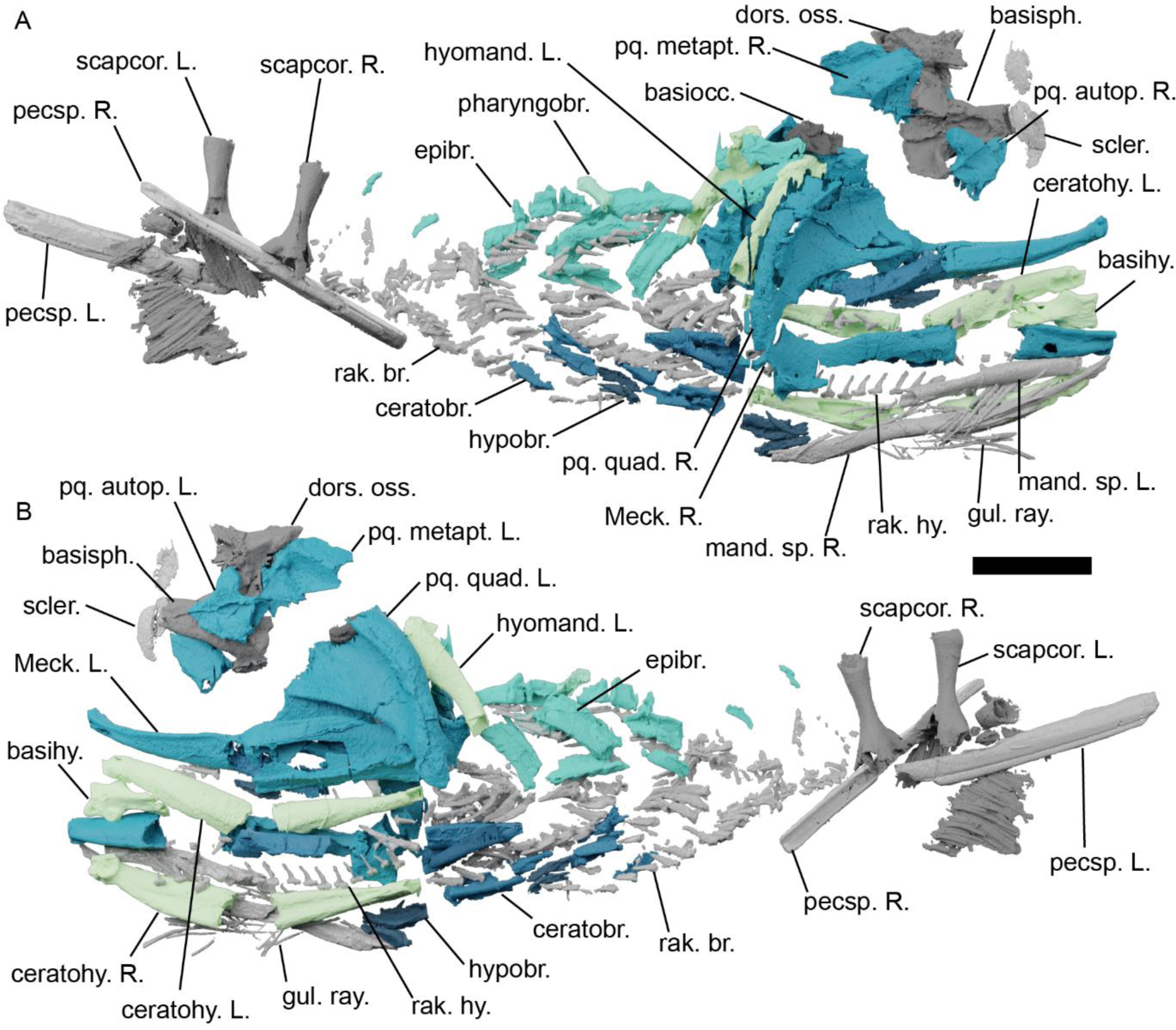
The head skeleton of *Acanthodes confusus* MNHN SAA20, visualised using computed tomography. A, in right lateral view. B, in left lateral view. Abbreviations: basihy., basihyal; basiocc., basioccipital; basisph., basisphenoid; ceratobr., ceratobranchials; ceratohy., ceratohyal; dors. oss., dorsal ossification of neurocranium; epibr,eprbranchial; gul. ray., gular rays; hyomand., hyomandibula; hypobr., hypobranchials; L., left; mand. sp., mandibular splint; Meck., Meckel’s cartilage; pharyngobr., pharyngobranchials; pecsp., pectoral fin spine; pq. autop., palatoquadrate autopalatine; pq. metapt., palatoquadrate metapterygoid; pq. quad., palatoquadrate quadrate; R., right; rak.br., branchial rakers; rak. hy., hyoid rakers; scapcor., scapulocoracoids; scler, sclerotic ring. Colour scheme: blue-greens, elements of visceral endoskeleton with mandibular arch, hyoid arch, hypobranchials, ceratobranchials, epibranchials, and pharyngobranchials coloured independently; dark grey, cranial and pectoral endoskeleton; light grey, elements of the dermal skeleton. Scale bar = 20 mm.

**Figure 4.**
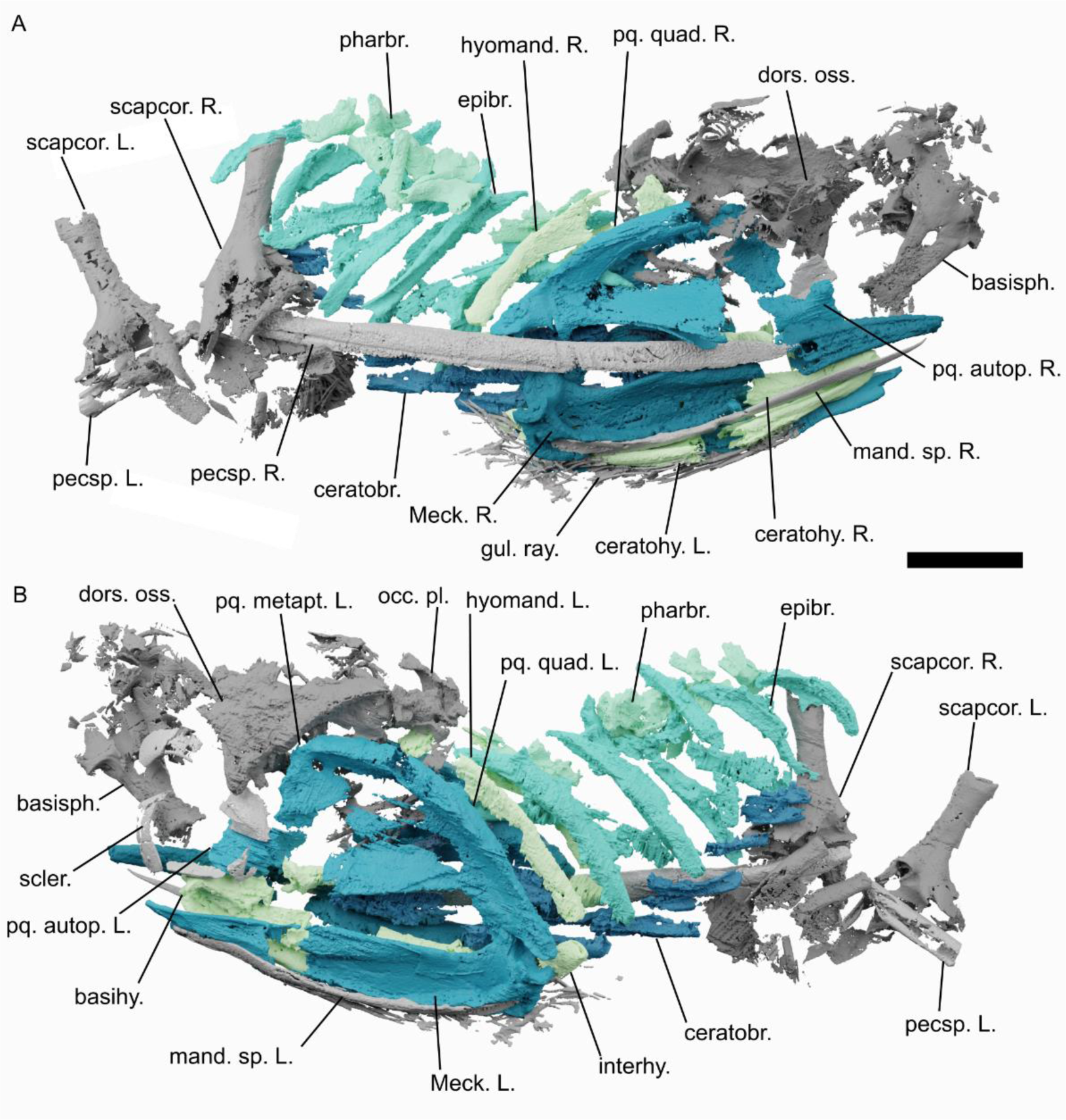
The head skeleton of *Acanthodes confusus* MNHN SAA24, visualised using computed tomography. A, in right lateral view. B, in left lateral view. Abbreviations: basihy., basihyal; basiocc., basioccipital; basisph., basisphenoid; ceratobr., ceratobranchials; ceratohy., ceratohyal; dors. oss., dorsal ossification of neurocranium; epibr,eprbranchial; gul. ray., gular rays; hyomand., hyomandibula; hypobr., hypobranchials; interhy., interhyal; L., left; mand. sp., mandibular splint; Meck., Meckel’s cartilage; occ. pl., occipital plate; pharyngobr., pharyngobranchials; pecsp., pectoral fin spine; pq. autop., palatoquadrate autopalatine; pq. metapt., palatoquadrate metapterygoid; pq. quad., palatoquadrate quadrate; R., right; scapcor., scapulocoracoids; scler, sclerotic ring. Colour scheme: blue-greens, elements of visceral endoskeleton with mandibular arch, hyoid arch, hypobranchials, ceratobranchials, epibranchials, and pharyngobranchials coloured independently; dark grey, cranial and pectoral endoskeleton; light grey, elements of the dermal skeleton. Scale bar = 20 mm.

### 3D Data Acquisition

We scanned the specimens using micro-computed tomography (μCT) using a Baker Hughes Digital Solutions v|tome|x 240 L μCT scanner at the MNHN AST-RX platform. SAA20 was scanned with a 1 mm Cu and a 0.5 Sn filter at 245 kV, resulting in a voxel size of 59 μm. SAA21 was scanned with a 0.3 mm Cu filter at 245 kV, resulting in a voxel size of 64 μm. SAA24 was scanned with a 1 mm Cu filter at 225 kV, resulting in a voxel size of 56 μm. We processed the tomographic data using Materialise Mimics 19.0 to create 3D models: models were manually segmented using ‘Edit Masks’ and ‘Multiple Slice Edit’ tools. We then rendered these models in the freeware Blender 3.30 (blender.org) to create images and videos

### Reconstruction and gape animation

We reconstructed the visceral skeleton of *Acanthodes* in Blender using models from the three specimens described here. We tidied the surface models by mending holes and smoothing surfaces. Unmineralised sections of bones were interpolated using the Blender sculpting tool. Models were then rearranged into an approximation of life position. To investigate the jaw opening mechanism we attached an armature to the jaws, with separate bones between the two palatoquadrate-skull articulations, between the upper articulation, and along Meckel’s cartilage. This was animated by making the palatoquadrates swing outwards around the axis of their articulation with the braincase. Meanwhile the Meckel’s cartilages were pulled ventrally relative to their articulation with the palatoquadrates.

## Results

All three specimens have been laterally flattened. Between them they preserve most of the known endoskeleton of *Acanthodes* (Figs 2-4) including includes the jaws, hyoid arch, branchial skeleton, braincase, and shoulder girdle. Here we concentrate on the visceral skeleton: the braincase and shoulder girdle will be the focus of future descriptions. Generally speaking the details of the visceral skeleton of *Acanthodes* matches the detailed accounts of Miles (1964, 1965, 1968, 1973). In this account we generally follow Miles’ terminology; a comparison of the terminology we use to that used by previous authors is given in Supplementary Table 1.

All endoskeletal elements in *A. confusus* comprise a heavily mineralised outer shell surrounding an internal space (Fig. 5). Although our scan data is not sufficiently high resolution to show the histology of the outer tissue, Ørvig (Ørvig, 1951) interpreted the same tissue as perichondral bone in thin sections of acanthodians including *Acanthodes* specimens from Lebach on the basis of the tissue having fusiform cell spaces with canaliculi and Sharpey’s fibre attachments, albeit lacking lamination or vascular canals. The thickness of this tissue is variable through the skeleton, for example being thicker in the mandibular arches than in the branchial arches, particularly around the mandibular articulation (Fig. 5c). This presumably reflects the function of different elements analogously to varying thicknesses in the prismatic tesselate calcified cartilage in extant and extinct chondrichthyans (Maisey *et al*., 2020). *Acanthodes* is the only known articulated total-group chondrichthyan in which the endoskeleton is extensively ossified, and the relationship of its perichondral bony tissue to varied cartilages in other stem-chondrichthyans is as yet unclear (Burrow *et al*., 2018, 2020; Maisey *et al*., 2020; Burrow & Blaauwen, 2021).

**Figure 5.**
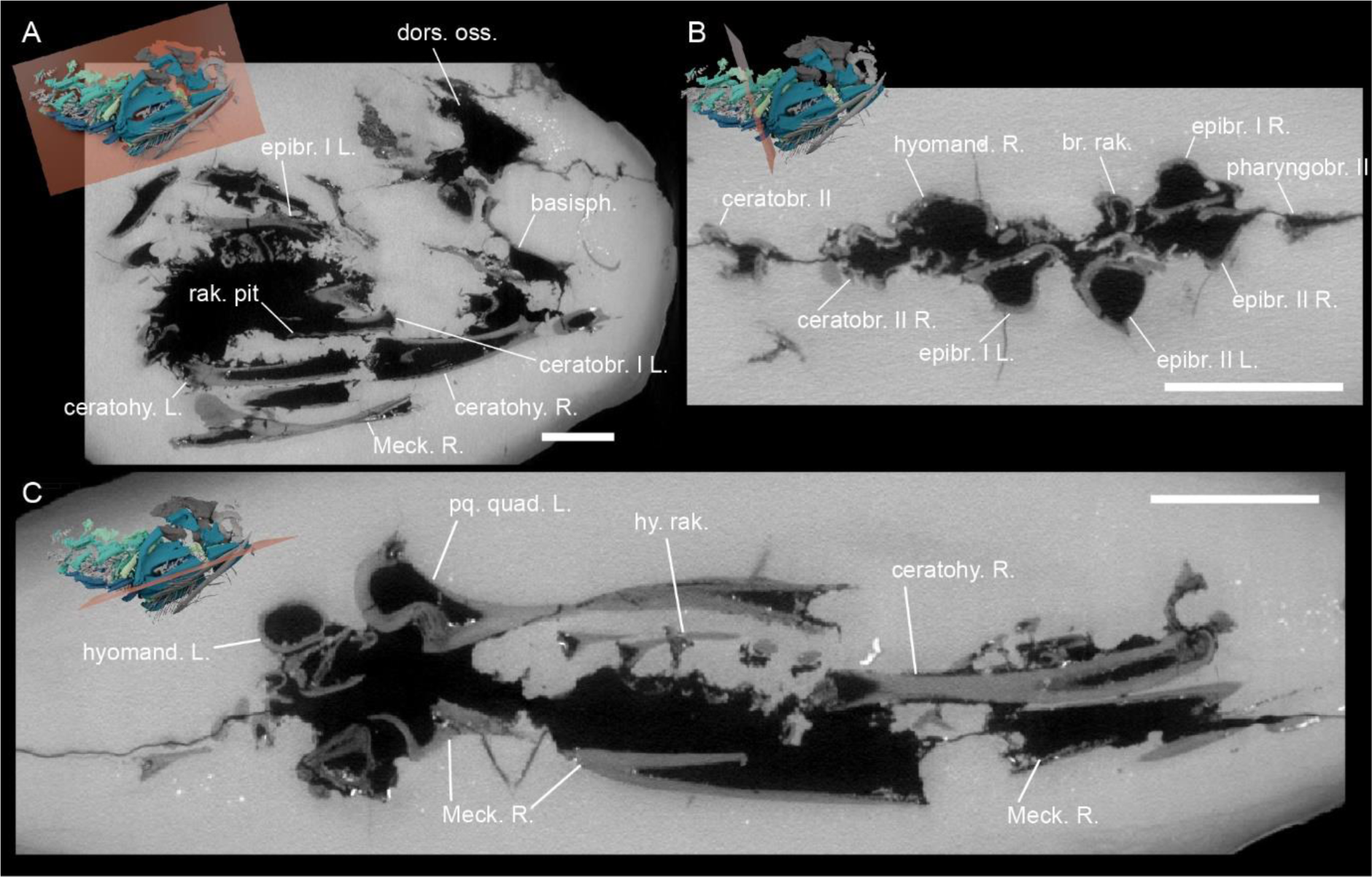
Sections through tomograms from the scan data for SAA21. A, sagittal section. B, transverse section. C, coronal section. Abbreviations: basisph., basisphenoid; br. rak., branchial rakers; ceratobr., ceratobranchials; ceratohy., ceratohyal; dors. oss., dorsal ossification of neurocranium; epibr,eprbranchial; hyomand., hyomandibula; hy. Rak., hyoid rakers; L., left; Meck., Meckel’s cartilage; pharyngobr., pharyngobranchials; pq. metapt., palatoquadrate metapterygoid; R., right, rak. pit, pit for pharyngeal raker. Red plane indicates plane of bisection. Scale bar = 10 mm.

### Mandibular arch

The palatoquadrates are ossified in three parts – metapterygoid, autopalatine, and quadrate (Figs 2-4, 6A,B,G) – with a tall otic process and pronounced extrapalatoquadrate ridge. In cross-section the commissural lamina of the palatoquadrate is laterally convex forming a large attachment surface for the mandibular adductor muscle, on the medial face this forms a medial angle (Fig. 6B). The palatoquadrate is cleaver-shaped in profile similarly to other stem-group chondrichthyans is seen in *Ptomacanthus, Climatius, Cheiracanthus,Ischnacanthus, Mesacanthus* (Watson, 1937; Miles, 1973b; Brazeau, 2012; Burrow *et al*., 2015, 2018, 2022). A foramen in the metapterygoid seems most plausibly to have carried the mandibular ramus (V_3_) of the trigeminal nerve (Miles, 1964). Each palatoquadrate articulates with the braincase at three points (Miles, 1968): by two condyles on the metapterygoid which articulate with the postorbital part of the braincase and a rounded basal process on the autopalatine which articulates with the basisphenoid (Figs 2, 6). The anterior edge of the auropalatine is finished, with no evidence for an anteriorly extending palatine commissure as interpreted by Jarvik (1977).

**Figure 6.**
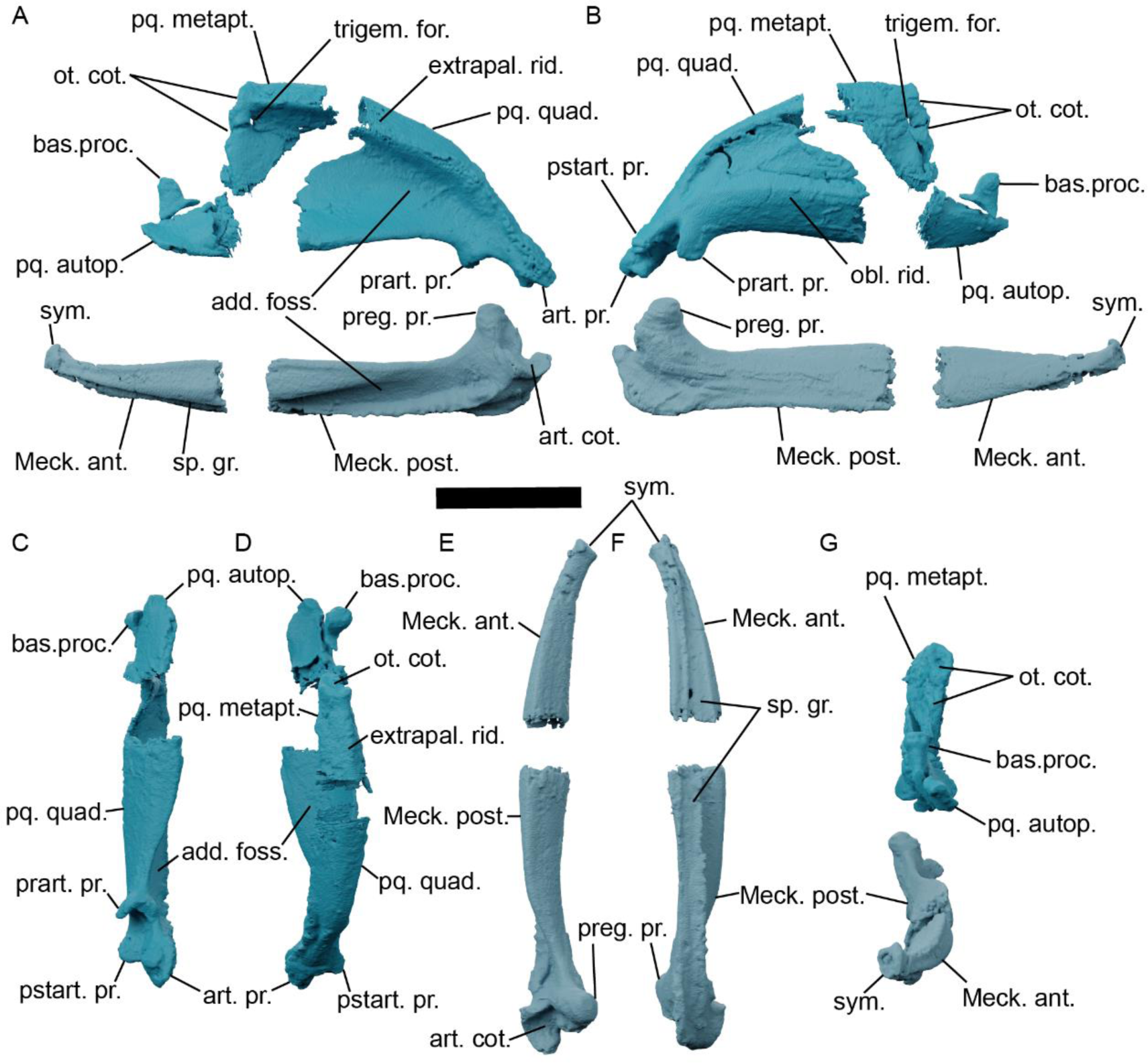
The mandibular arch of *Acanthodes confusus* MNHN SAA21. A, left Meckel’s cartilage and palatoquadrate in lateral view. B, left Meckel’s cartilage and palatoquadrate in medial view. C, D, left palatoquadrate in (C) ventral, and (D) dorsal view. E,F, left Meckel’s cartilage in (E) dorsal, and (F) ventral view. G, left Meckel’s cartilage and palatoquadrate in anterior view. Abbreviations: add. foss., fossa for the adductor muscle; art. cot., articular cotylus; art. pr., articular process; bas. proc., basal process; extrapal. rid., extrapalatoquadrate ridge; Meck. ant., anterior mineralisation of Meckel’s cartilage; Meck. post., posterior mineralisation of Meckel’s cartilage; obl. rid., oblique ridge; ot. cot., cotylus for articulation with otic region; pq. autop., palatoquadrate autopalatine; pq. metapt., palatoquadrate metapterygoid; pq. quad., palatoquadrate quadrate; prart. pr., prearticular process; preg. pr., preglenoid process; sp. gr., mandibular splint groove; sym. expanded mandibular symphysis; trigem. for., foramen for the mandibular ramus of the trigeminal nerve (V_3_). Scale bar = 20 mm.

The Meckel’s cartilage is ossified in two parts (Miles, 1968, 1973a), and in overall shape tapers anteriorly and curves medially (Fig. 2-4, 6C-F). A large lateral fossa provided an insertion site for the mandibular adductor while a ventrolateral groove would have carried the dermal mandibular splint, which matches the description of Dearden & Giles (2021). The mandibles are separate, but the expanded anterior tip suggests a well-developed symphysis, which may be a character uniting some stem-group chondrichthyans (Dearden & Giles, 2021).

The palatoquadrate articulates with the mandible via a chondrichthyan-like double articulation with an articular process and glenoid fossa on the palatoquadrate articulating with a preglenoid process and articular fossa on the mandible (Miles, 1968) (Fig. 6). The preglenoid process is notably more rounded than as reconstructed by Miles (1968). This bears comparison to other chondrichthyans (Miles, 1973a) including Palaeozoic forms such as *Gogoselachus* (Long *et al*., 2015) and *Gladbachus* (Coates *et al*., 2018). However a pronounced retroarticular flange as in *Gogoselachus* and *Tristychius* is absent (Long *et al*., 2015; Coates *et al*., 2019; Frey *et al*., 2020). When closed the jaws are tilted slightly dorsally, as reconstructed by Brazeau & de Winter (2015).

### Hyoid arch

The hyoid arch comprises paired hyomandibulae, interhyals, ceratohyals, and a basihyal (Figs 2-4,7,8). The hyomandibula comprises a posterior and poorly mineralised anterior ossification (Fig. 8A,B). We are unable to confirm its articulation relative to the jugular vein groove but we find nothing to suggest the interpretation of its articulation by Brazeau and de Winter (2015) is inaccurate. The end of the hyomandibula contacting the neurocranium is laterally compressed in cross-section, grading into a circular cross-section posteriorly, and the medial face of the hyomandibula is split by a marked, longitudinal ridge, which separates it into a dorsomesial and ventromesial faces (Fig. 7F)(Miles, 1968). The laterally flattened, curved, and posteriorly tapered shape of the hyomandibula is more comparable to that of the actinopterygian *Mimipiscis* (Gardiner, 1984), symmoriiforms *Ozarcus* (Pradel *et al*., 2014) and *Ferromirum* (Frey *et al*., 2020) and cladoselachians (Maisey, 1989) than to the stem-group chondrichthyan *Gladbachus* (Coates *et al*., 2018) or to the stouter hyomandibulae of crown-group chondrichthyans such as xenacanths or hybodonts (Hotton, 1952; Maisey, 1987).

**Figure 7.**
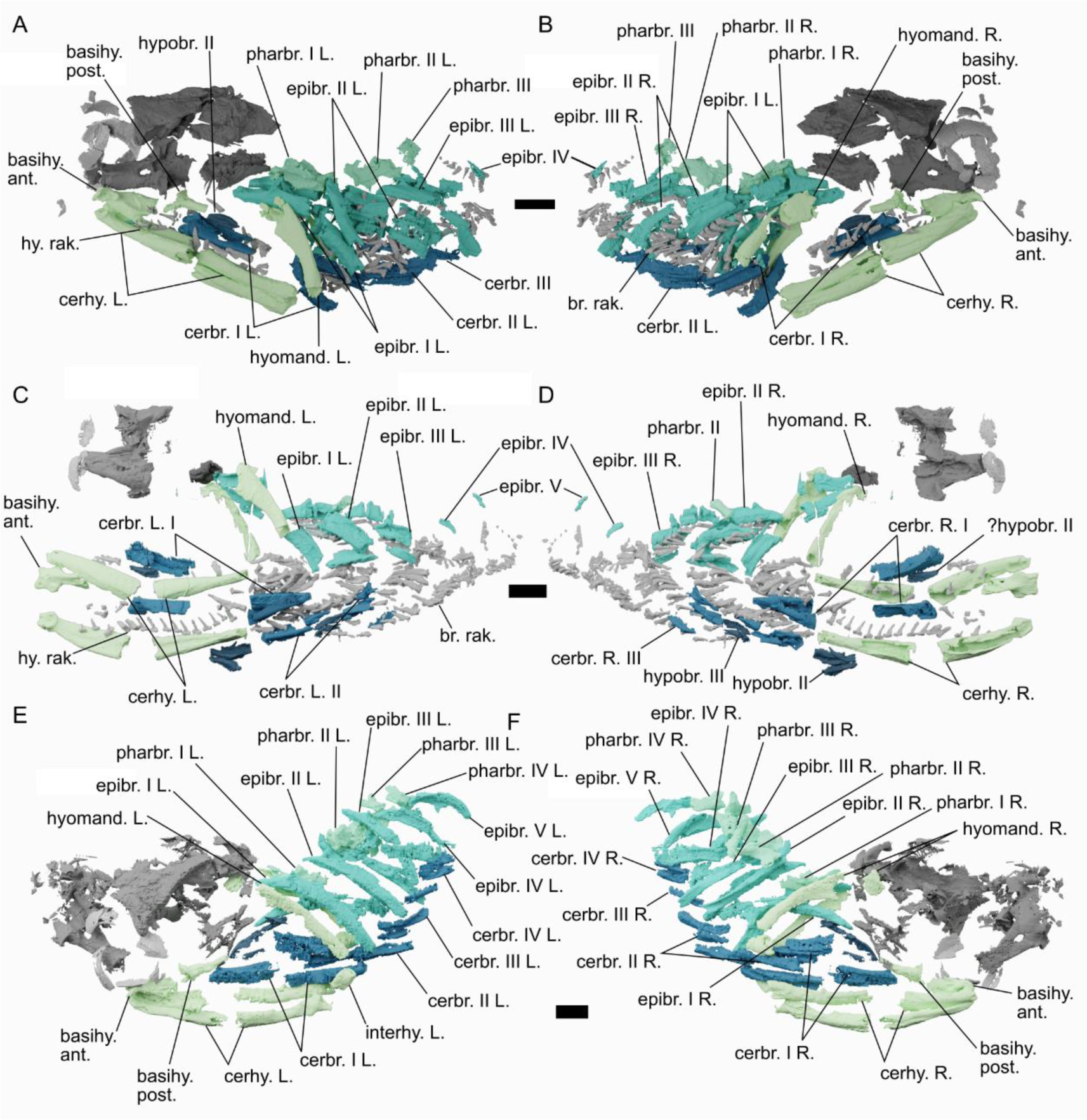
Overview of *Acanthodes confusus* hyoid and branchial skeleton. A,B, SAA21 with mandibular skeleton removed in (A) left lateral and (B) right lateral view. C, D. SAA20 with mandibular and pectoral skeleton removed in (C) left lateral and (D) right lateral view. E, F, SAA24 with mandibular and pectoral skeleton removed in (E) left lateral and (F) right lateral view. Abbreviations: basihy., basihyal; ceratobr., ceratobranchials; cerhy., ceratohyal; epibr, epibranchial; hyomand., hyomandibula; interhy., interhyal; L., left; Meck., Meckel’s cartilage; pharbr., pharyngobranchials; pq. autop., palatoquadrate autopalatine; pq. metapt., palatoquadrate metapterygoid; pq. quad., palatoquadrate quadrate; R., right; rak.br., branchial rakers; rak. hy., hyoid rakers. Colour scheme: blue-greens, elements of visceral endoskeleton with mandibular arch, hyoid arch, hypobranchials, ceratobranchials, epibranchials, and pharyngobranchials coloured independently; dark grey, cranial and pectoral endoskeleton; light grey, elements of the dermal skeleton. Scale bars = 10 mm.

**Figure 8.**
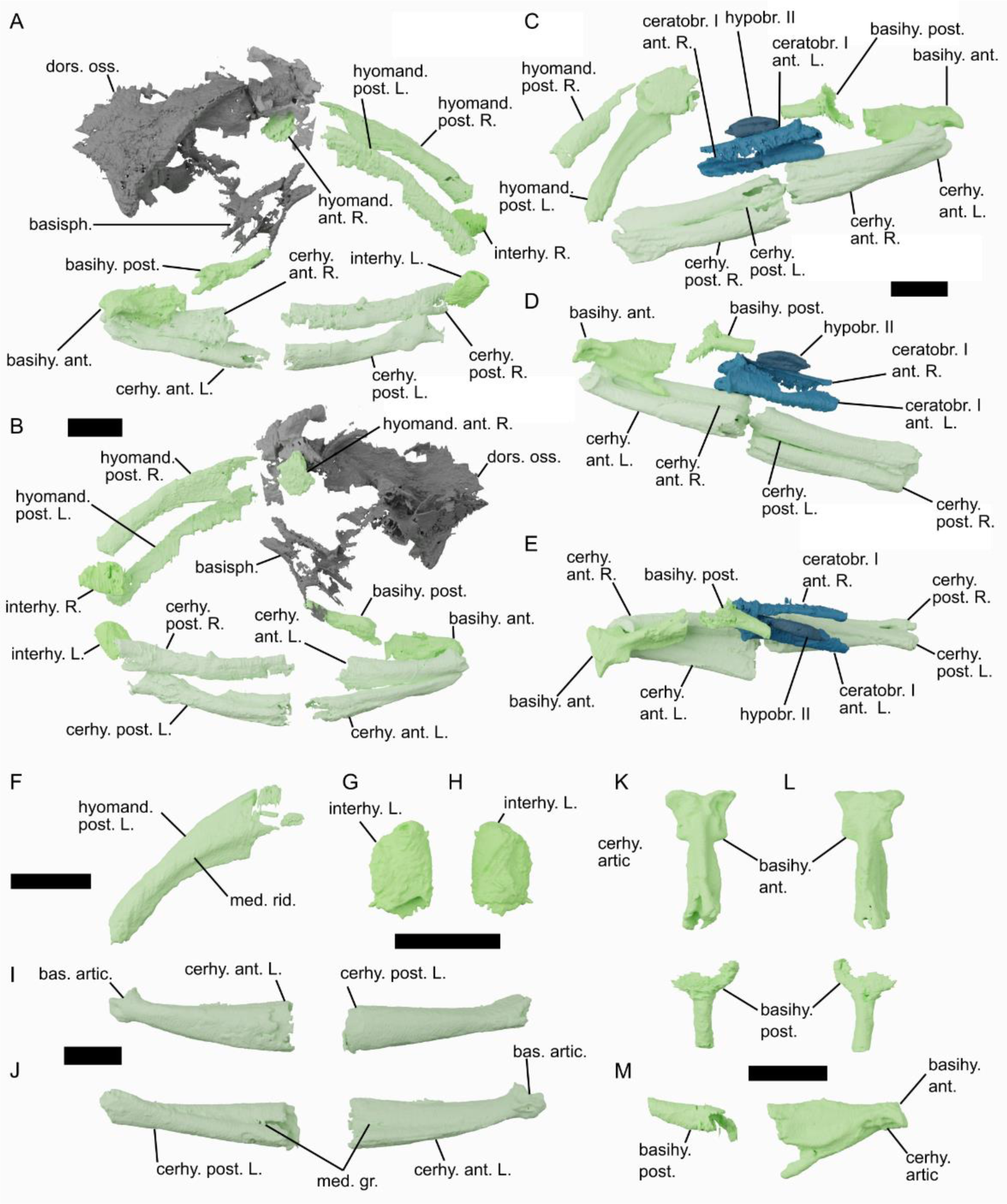
The hyoid skeleton of *Acanthodes confusus*. A,B, the hyoid skeleton of SAA24 in (A) left lateral, and (B) right lateral view. C-E, the hyoid skeleton of SAA21 in (C) right lateral, (D) left lateral, and (E) dorsal view. F, the posterior mineralisation of the left hyomandibula in SAA21. G, H, the left interhyal of SAA24 in (G) lateral, and (H) medial view. I,J, the left ceratohyal of SAA21 in (I) lateral, and (J) medial view. K-M, the basihyal of SAA21 in (K) ventral, (L) dorsal, and (M) right lateral view. Abbreviations: ant., anterior ossification; artic., articulation; basihy., basihyal; basisph., basisphenoid; cerhy., ceratohyal; dors. oss., dorsal ossification of neurocranium; L., left; hyomand., hyomandibula; hypobr., hypobranchials; med. gr., medial groove; med. rid., medial ridge; post., posterior ossification; R., right.; I-V, branchial arches I-V. Scale bar = 10 mm

An interhyal is present between the hyomandibula and ceratohyal of *Acanthodes* (Figs 4, 7,8A,B,G,H). The interhyal is subrectangular and laterally flattened, with gently convex dorsal and ventral surfaces. Of the three specimens scanned, interhyals are only preserved in SAA24 (Fig. 4) and before that were only known from a single mouldic specimen (MFN MB 23) (Miles, 1965, 1973a; Davis *et al*., 2012), suggesting that whether or not they ossified was variable in different individuals. As Miles (1973) highlighted it is unclear whether this “interhyal” is homologous to the interhyal, stylohyal, or symplectic in osteichthyans (Patterson, 1982; Véran, 1988). Whether or not these are homologous, additional elements of the hyoid skeleton are not found in this position in *Gladbachus* (Coates *et al*., 2018), or in other articulated chondrichthyans (Pradel *et al*., 2014; Frey *et al*., 2020; Klug *et al*., 2023).

The ceratohyals are each ossified in two parts with no hypohyals present (Figs 2-4, 7,8A-E,I,J) consistent with the reconstructions of Miles (1965, 1973) and Gardiner (1984). The posterior end is laterally flattened and lacks the lateral fossa seen in some early chondrichthyans (Coates *et al*., 2018). It also lacks the sharp dorsal angle at the posterior end of the ceratohyals of *Ferromirum* (Frey *et al*., 2020), *Phoebodus* (Frey *et al*., 2019), and *Maghriboselache* (Klug *et al*., 2023). The elements profile bulges in the middle, narrowing again in its anterior half. The anterior end of the ceratohyal is not spatulate anteriorly like some Palaeozoic chondrichthyans such as *Phoebodus* (Frey *et al*., 2019), instead pinching in and expanding out to form the articulation with the basihyal (Miles, 1968). A longitudinal groove runs along the mesial surface of the element which may be homologous a similar groove on the ceratohyal of *Gydoselache* (Maisey *et al*., 2019).

The component forming the ventral floor of the pharyngeal skeleton is formed from two mineralisations (Figs 2-4, 7,8A-E,K-M). This has variously been termed a basibranchial or basihyal; here we use the latter as its articulation with the hyoid and first branchial arch is more comparable to the arrangement in living chondrichthyan basihyals than to osteichthyan basibranchials. The anterior mineralisation has an “hammerhead” anterior end with ventrally-oriented articulation surfaces for the ceratohyals. In its posterior half this becomes taller with deep ventral attachment surfaces for the coracohyoid and coracobranchial musculature, with an unfinished posterior face. The posterior basihyal mineralisation is flat with a posterior tail, its unfinished anterior face suggests that it was joined to the anterior mineralisation of the basihyal by cartilage. There are no obvious articulation surfaces on the posterior part of the basihyal, but based on their preserved position it seems likely to have articulated with the first and second branchial arch as reconstructed by Miles (Miles, 1973a). This tall, narrow basihyal is dissimilar from the broad flat basihyals of other known stem-group chondrichthyans (Brazeau, 2012; Coates *et al*., 2018), although a tapering posterior end is common in basibranchial copulae (Coates *et al*., 2018; Dearden *et al*., 2021). A possible exception is in *Halimacanthodes,* where an element identified as an ?autopalatine (or possibly a basibranchial), could be the front of the basihyal in lateral profile (Burrow *et al*., 2012). We find no evidence for a chain of basibranchial elements as given in some reconstructions (Fig. 1A-C): this aspect of the reconstruction appears to be based on the holotype of *A. confusus* (Heidtke, 2011, fig. 8) in which preserved hypobranchials may be giving the impression of there being a chain of basibranchials.

### Branchial arches

Five branchial arches are preserved which extend well-posterior to the braincase (Figs 2-4, 6,8). It is unclear from our data whether they articulated with the basioccipital, as reconstructed by Miles (1973a), but in SAA21 they overlap with the underside of the braincase anteriorly (Dearden *et al*., 2019; Frey *et al*., 2020). The branchial arches comprise hypo-, cerato-, epi-, and pharyngobranchials. There is no evidence for accessory elements like in some osteichthyans and in the symmoriiform *Ozarcus* (Pradel *et al*., 2014).

Four pairs of ceratobranchials are preservd, a fifth ceratobranchial is absent although rakers on the ventral part of the fifth arch in SAA20 (fig. 7C) indicate that one was probably present but unossified. The ceratobranchials all have a similar overall structure, with a ventral groove for the afferent branchial artery and indentations on their dorsal surface for the branchial rakers (Fig.9L). Ceratobranchial I and II are segmented into anterior and posterior sections, and ceratobranchial I has a pronounced articular facet on its anterior end. More posterior ceratobranchials become progressively shorter and more pronouncedly curved until the fourth pair is almost as broad as they are long (Fig. 7E,F). The posteriormost ceratobranchial is not enlarged as in *Gladbachus* and the stem-group gnathostome *Paraplesiobatis* (Brazeau *et al*., 2017; Coates *et al*., 2018), but similarly to them is more flattened relative to the others.

**Figure 9.**
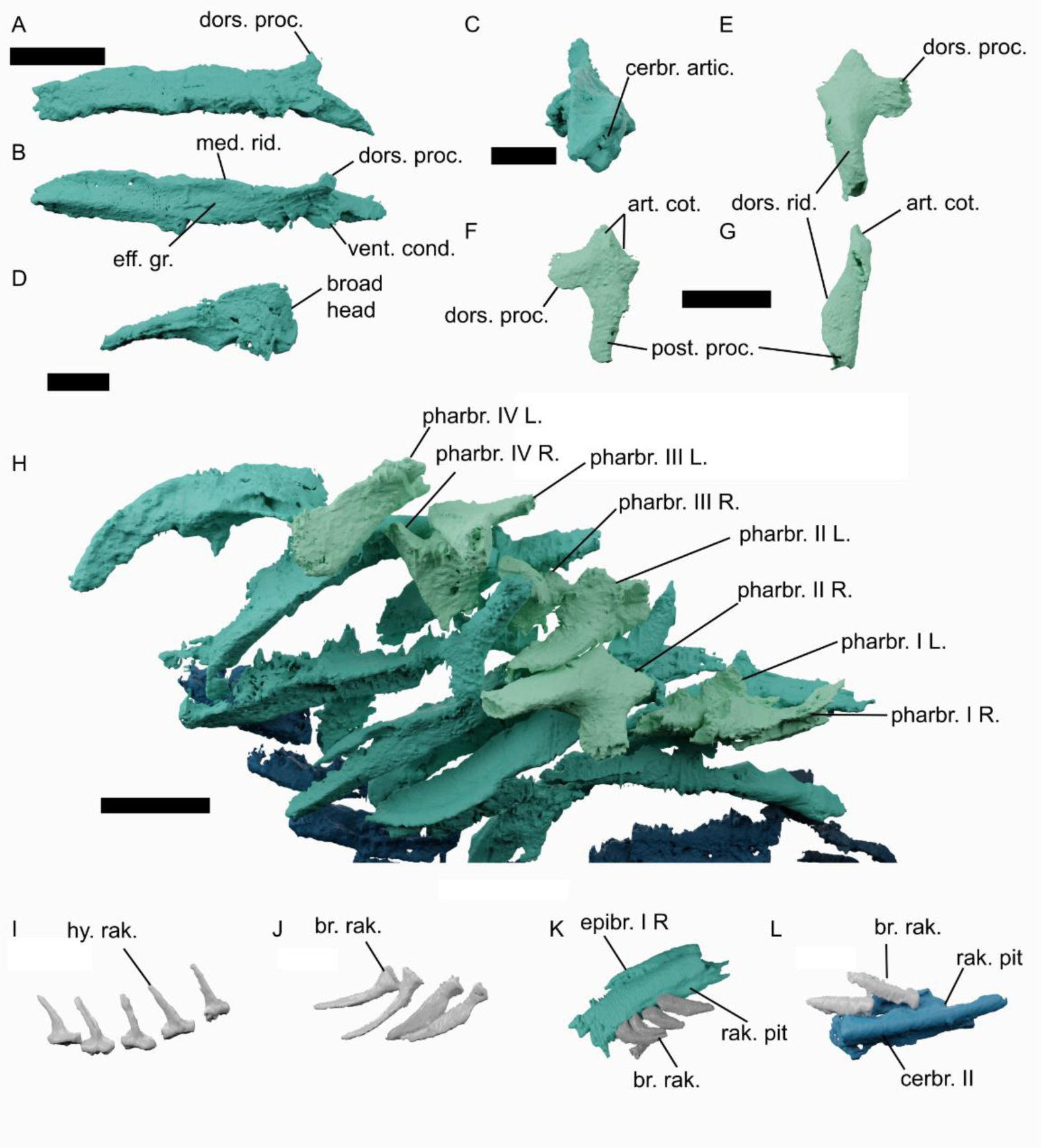
The branchial skeleton of *Acanthodes confusus*. A-C, the right epibranchial II of SAA24 in (A) lateral, (B) dorsal, and (C) posterior view. D, the left fifth epibranchial of SAA24 in dorsal view. E-G, the right pharyngobranchials II of SAA24 in (E) dorsal, (F) ventral, and (G) lateral view. H, articulated dorsal branchial skeleton of SAA24 in right lateral view with pharyngobranchials labelled. I, hyoid rakers from the ceratohyal of SAA20. J, branchial rakers from the epibranchial of SAA20. K, branchial rakers in articulation with the epibranchial of SAA21. L, branchial rakers in articulation with the ceratobranchial of SAA21. Abbreviations: artic., articulation; art. cot. articular cotylus; cerbr., ceratobranchial; dors. proc., dorsal process; dors. rid., dorsal ridge; epibr., epibranchial; L, left; med. rid., medial ridge; pharbr., pharyngobranchial; post. proc., posterior process; R, right; rak.br., branchial rakers; rak. hy., hyoid rakers; rak. pit, pit for branchial rakers; vent. cond, ventral condyle; I-V, branchial arches I-V Scale bar = 10 mm except in panel c where scale bar = 2.5 mm.

We can confirm there are hypobranchials on at least the second and third branchial arches based on our scan data (Fig. 7). We can see no hypobranchial IV, although a cast figured by Miles (1973, plate 7) seems to show a hypobranchial in this position quite clearly and this may be either lost or unmineralized in the specimens studied here. Hypobranchials are short and curved, with a lateral groove, and are oriented anteriorly. Ceratobranchial I has a very well-developed condyle at its anterior tip and is preserved in close association with the basihyal in SAA21 (Fig. 8C,D), suggesting a direct connection between the two. There is no evidence for a fifth hypobranchial.

Five pairs of epibranchial are present, one in each arch (Figs 7, 9). The epibranchials are gently curved, with a dorsolateral groove for the efferent branchial artery (Fig. 9A,B). This groove is bordered medially by a ridge corresponding to the posterior flange described by Coates *et al*. (Coates *et al*., 2018) that at the distal end of the ceratobranchial develops into a dorsal process (Miles, 1968). Like the ceratobranchials the surface facing into the branchial chamber is pitted to carry the gill rakers (Fig. 9K). The proximal end of the epibranchials has an articular surface for the ceratobranchial (Fig. 9C). The epibranchials have the same overall form, but become progressively shorter posteriorly (Figs 7, 9H). The posteriormost (fifth) epibranchials are different in shape, with broad heads (Figs 7, 9D, H). Although the epibranchials in SAA20 and SAA21 appear to be segmented, in SAA24 they are ossified into a single structure (Fig. 4A,C).

There are four pairs of pharyngobranchials, best preserved in SAA24 (Fig. 4C, 7C, 9E-H). As in previous accounts there is no evidence for separate supra and infrapharyngobranchials as in *Ozarcus* and osteichthyans (Gardiner, 1984; Pradel *et al*., 2014). These closely match the anatomy and orientation of the pharyngobranchials in *Halimacanthodes* (Burrow *et al*., 2012). The interpretation of the articulation with the epibranchial given by Miles, also seems to be accurate, with the right epibranchial II in SAA24 preserved in this position (Fig. 8H), albeit with the result that they are oriented as reconstructed by Jarvik (1977, fig. 8). As he surmised this results in the pharyngobranchials oriented ventro-medially, presumable leaving the surface between the posterior process and dorsal surface as an attachment area for the *m. interpharyngobranchialis* (as figured by Jarvik, 1977). There is no evidence for a pharyngobranchial anterior to the first epibranchial. Based on this we consider the interpretation of Nelson, Miles, and Jarvik (Nelson, 1968; Miles, 1973a; Jarvik, 1977) to be accurate, in that these represent the first four, posteriorly-oriented pharyngobranchials. Unlike living elasmobranchs, but like *Gladbachus* and *Ptomacanthus* (Coates *et al*., 2018; Dearden *et al*., 2019)the posteriormost epibranchials and pharyngobranchials are not fused into a single complex element.

### Hyoid and branchial rakers

Both branchial arches and the hyoid arch carry rows of small rakers. Our data is not sufficiently high resolution to show their histology, but these elements are separate from the endoskeletal arches, with distinct bases and crowns, and are ornamented (Zidek, 1985) so we consider it probable they are dermal rakers (Miles, 1968) rather than endoskeletal projections from the pharyngeal arches (Ørvig, 1973; Jarvik, 1977). On the basis of SAA21 and SAA20 (Figs 2,3) there is a single row of antero-medially directed rakers on each branchial arch, although more than one row on each arch has been reported, which may vary through ontogeny (Reis, 1896; Watson, 1937; Miles, 1968). The ceratohyals carry a single row of rakers (Figs 2,3,7). These are distinctly different in morphology from the branchial rakers: the hyoid rakers are shorter and straight with flat bases (Fig 9I) whereas those on the branchial arches are longer, slightly curved, with a distinct neck and a concave base (Fig 9J). The largest hyoid rakers are in the middle of the ceratohyal, whereas the largest branchial rakers are in the centre of each gill arch (i.e. near the junction between the cerato- and epibranchial (Fig 2,3,7). We find no evidence for rakers on the hyomandibula in our data, which conflicts with reports from previous descriptions (Reis, 1896; Watson, 1937; Miles, 1968)

### Functional morphology

Our 3D reconstruction confirms that the reconstruction of Miles with three points of articulation with the neurocranium is plausible (Fig. 10). The effect of animating this is that the palatoquadrates swing laterally as he predicted. The double articulation of the Meckel’s cartilage and the palatoquadrate means that the mandible could only lower vertically relative to the palatoquadrate, meaning that the relative angles of the left and right mandibles change during jaw opening. This is perhaps the reason for the expanded symphyseal tip on the mandible, to articulate with connective tissue that allows this movement. Miles also suggested that the hyoid arch was the mechanism for pushing the jaws outwards: the very well developed articulatory surfaces between the ceratohyals and basihyal and on the first ceratobranchials, as well the large potential attachment surfaces on the basihyal suggest that the basibranchial may have been functionally important.

**Figure 10.**
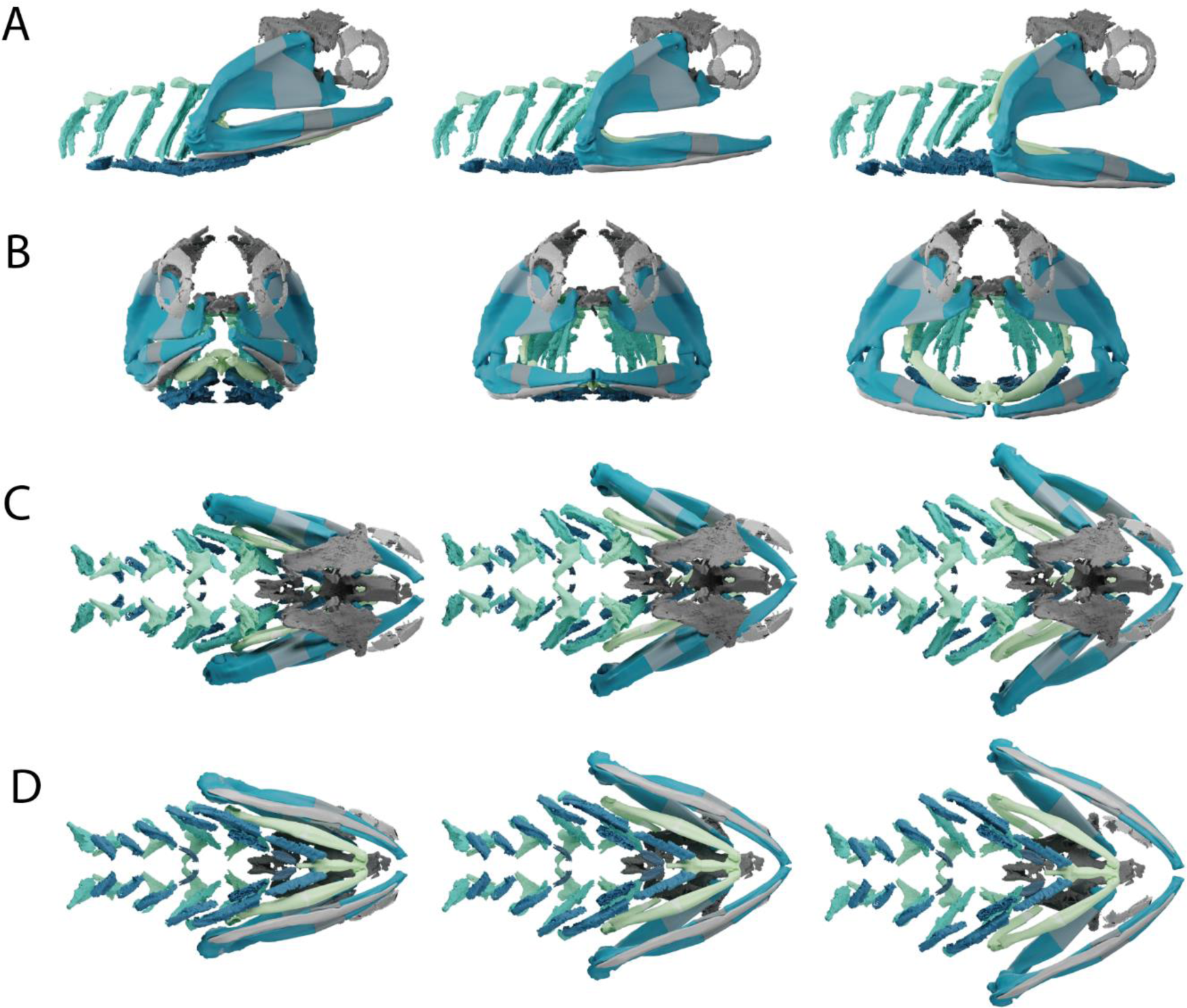
Reconstruction of *Acanthodes confusus* based on a composite of the material described here, animated to show mouth opening from left to right. A in right lateral view, B in anterior view, C in dorsal view, D in ventral view.

## Discussion

### Pharyngeal skeletal patterning in *Acanthodes*

Several reconstructions of the pharyngeal endoskeleton of *Acanthodes confusus* have been made, based exclusively on casts based on fossils from Lebach (Fig. 1). Early reconstructions were made by Reis (1890, 1894, 1895, 1896) and then by Dean (1907). Watson (1937) then reconstructed the skeleton and used it as the basis for his argument that acanthodians were aphetohyoidean. Renewed interest in the late 20^th^ century then led to reconstructions by Miles (1964, 1968, 1973), Nelson (1968), Jarvik (1977) and Gardiner (1984). Since then no new information on the branchial skeleton has been published (although it has been discussed (Coates *et al*., 2018; Dearden *et al*., 2019) and Davis (2012) and Brazeau and de Winter (2015) discussed the articulation of the hyomandibula with the braincase. Disagreement between these reconstructions centres around the presence/absence of hyoid components (interhyal, pharyngohyal, and hypohyal), the structure of the basihyal/branchial skeleton, and the orientation of any hypobranchials and pharyngobranchials. Comparison between these reconstructions shows that much of this disagreement is caused by the segmentation of the pharyngeal skeleton (Fig. 1).

Our CT data allow us to arbitrate between previous reconstructions (Fig. 11) and place this anatomy on the context of recent advances in our understanding of Palaeozoic gnathostome pharyngeal anatomy. Generally speaking the most recent reconstructions by Miles (1964, 1968, 1973) Gardiner (1984), and Nelson (1968) are the most accurate, with them correctly identified which borders represent the unmineralised sections of elements. The hyoid arch of *Acanthodes* comprised a ceratohyal, interhyal, and hyomandibula (epihyal). The basibranchial skeleton comprised a single basihyal element which articulated with the ceratohyals and the first branchial arch. Hypobranchials are present on the second, third, and probably fourth arches, oriented anteriorly, and pharyngobranchials are present and oriented posteriorly. A single row of antero-medially oriented rakers is present on epibranchials and ceratobranchials, as well as on the ceratohyals.

**Figure 11.**
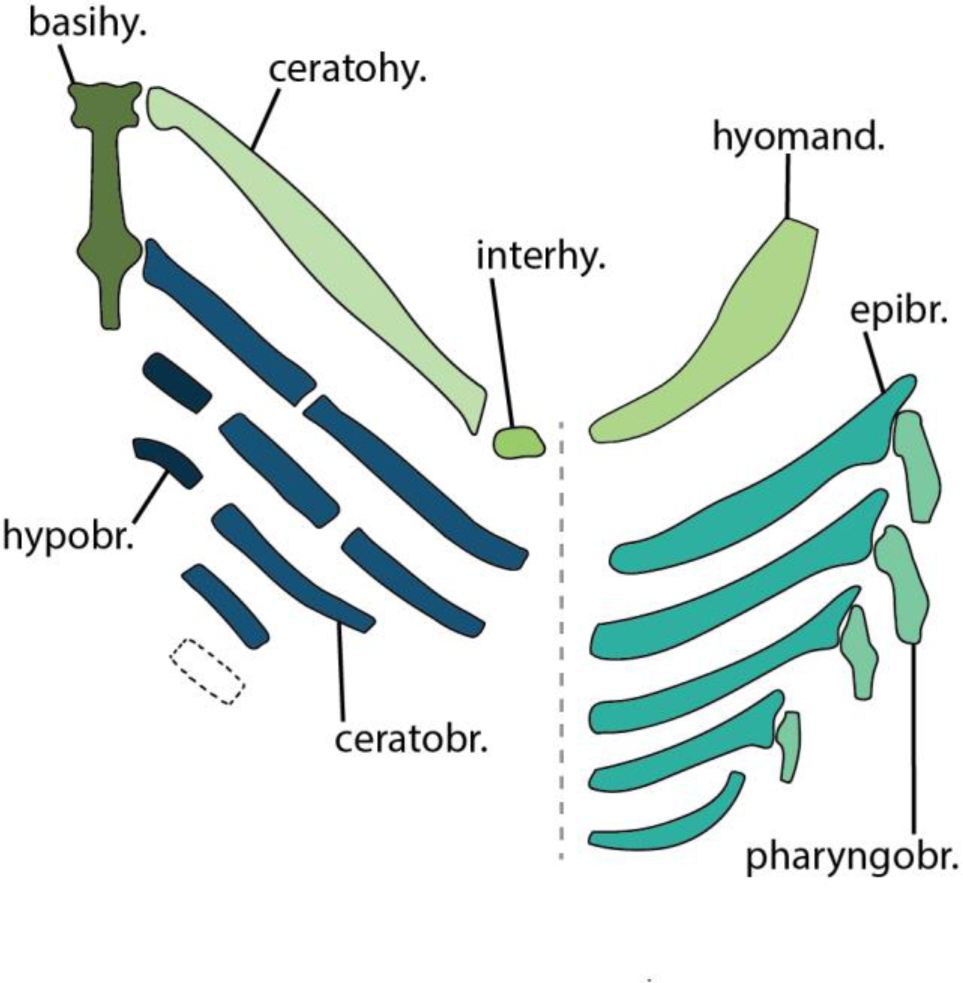
Reconstruction of the pharyngeal skeleton of *Acanthodes confuses* as described here. . Abbreviations: basihy., basihyal; cerbr., ceratobranchials; cerhy., ceratohyal;; epibr,epibranchial; hyomand., hyomandibula; interhy., interhyal; pharyngobr., pharyngobranchials. Matching colours denote serially homologous elements.

Phylogenetic characters based on the branchial skeleton are necessarily based on interpreting the presence/absence of small, often-poorly mineralized components of a delicate skeletal structure. The multiple specimens of *Acanthodes* that we scanned, as well as other acid-prepared specimens, show how this fact combined with taphonomy can alter the interpretation of character states. For example the interhyal is preserved in only one of the three specimens of *Acanthodes* we scanned, and in only one other specimen besides.

The epibranchials, are preserved as a single unit in SAA24 but segmented in the other two specimens. Finally a fifth ceratobranchial is implied by the presence of a fifth, ventral set of branchial rakers but is absent in all specimens. Thus caution should be taken in interpreting branchial characters based on presence/absence in a single specimen.

### Osteichthyan and chondrichthyan pharyngeal character states

The pharyngeal skeleton of *Acanthodes* displays a combination of osteichthyan-like and chondrichthyan-like characters. Past interpretations of the pharyngeal anatomy of *Acanthodes* have varied between chondrichthyan-like (Jarvik, 1977) and osteichthyan-like (Miles, 1973a) models. Recent studies of the neurocranial anatomy of *Acanthodes* have identified chondrichthyan-like characters such as a dorsal otic ridge (Davis *et al*., 2012) and a hyomandibular articulation ventral to the jugular vein (Brazeau & de Winter, 2015), which form a major part of evidence that *Acanthodes* is a stem-group chondrichthyan (Zhu *et al*., 2013; Coates *et al*., 2018; Dearden *et al*., 2019). However, the neurocranium also has severalosteichthyan-like traits, inferred to be gnathostome crown-group symplesiomorphies, including spiracular grooves on the basisphenoid and a tropibasic neurocranium (Friedman & Brazeau, 2010). Similarly, the branchial skeleton shows a combination of osteichthyan-like and chondrichthyan-like traits, although these are more difficult to polarize than cranial characters due to the dearth of information on the pharyngeal skeleton of stem-group gnathostomes (Carr *et al*., 2009; Brazeau *et al*., 2017). Here we discuss these based on the assumption that *Acanthodes* is indeed a stem-group chondrichthyan.

In overall aspect the pharyngeal skeleton is rather osteichthyan-like. In stem-group chondrichthyans *Ptomacanthus* and *Gladbachus*, as well as many extant elasmobranchs the basihyal, and thus the pharyngeal floor, is broad with widely spaced ceratohyals (Coates *et al*., 2018; Dearden *et al*., 2019, 2021). In *Acanthodes* the basihyal is narrow and the ceratohyals almost meet below the anterior end of the element. This bears comparison with the arrangement in Palaeozoic actinopterygians where the hyoid arch (albeit with hypohyals) articulate closely together with the antero-ventral surface of a narrow basibranchial (Gardiner, 1984; Giles *et al*., 2015). However, it is also comparable to the narrow-based pharynx of the symmoriiform *Ozarcus* (Pradel *et al*., 2014). Like *Ozarcus* and actinopterygians, *Acanthodes* has a narrow based neurocranium and it’s possible this is closely linked to cranial construction.

The construction of the hyoid arch in *Acanthodes* is similar in arrangement to that of other stem-group chondrichthyans in which it is known, like *Gladbachus* and *Ptomacanthus,* in that there is no separate hypohyal, no pharyngohyal, and a basihyal (Coates *et al*., 2018; Dearden *et al*., 2019). However, an interhyal is absent in both of these other taxa and is demonstrably absent in the vast majority of other chondrichthyans in which the hyoid skeleton is known such as *Gladbachus*, *Tristychius*, and *Ozarcus* (Pradel *et al*., 2014; Coates *et al*., 2018, 2019). A possible interhyal reported in a cladoselachian (Maisey, 1989) instead appears to represent the broken proximal end of the ceratohyal, split through the external fossa that is a common feature of Palaeozoic chondrichthyan ceratohyals (Coates *et al*., 2018). Amongst acanthodians an interhyal has also been reported in a specimen of *Ischnacanthus* (Brazeau & de Winter, 2015) and an “interhyal gap” (*interhyaler Spalt*) was identified in *Latviacanthus*, although without direct evidence an interhyal’s presence (Schultze & Zidek, 1982).

One or more skeletal elements between the ceratohyal and hyomandibula are present in osteichthyans, variously termed the interhyal, symplectic, and stylohyal, and have been cited as a character grouping them to the exclusion of chondrichthyans (Schaeffer, 1968; Patterson, 1982; Véran, 1988). Similarities were recently drawn between a cartilage in a stem-group gnathostome buchanosteid and the osteichthyan interhyal (Hu *et al*., 2017). Which, if any, of the osteichthyan hyoid elements the interhyal in *Acanthodes* is homologous with is unclear and ultimately obscured by a poor understanding of hyoid arch morphology in stem-group gnathostomes and Palaeozoic osteichthyans (Véran, 1988; Friedman & Brazeau, 2010). Regardless, the interhyal or symplectic of osteichthyans, the presence of a separate element between the ceratohyal and hyomandibula in *Acanthodes*, combined with its presence in a buchanosteid suggests that this may be a crown-group gnathostome symplesiomorphy.

Discussion of the pharyngobranchials in *Acanthodes* has in the past been framed in the context of contrasting states in living gnathostomes: anteriorly oriented infrapharyngobranchials and suprapharyngobranchials in osteichthyans vs posteriorly oriented pharyngobranchials (assumed to be homologous to infrapharyngobranchials) in chondrichthyans (Nelson, 1968; Miles, 1973a; Pradel *et al*., 2014). Our reconstruction confirms that *Acanthodes* was chondrichthyan-like in both respects, however since the last description of *Acanthodes* several more articulated chondrichthyan pharynxes have been described that show the distribution of character states is actually more complex than might be expected from modern taxa. *Gladbachus* has anteriorly oriented pharyngobranchials (but no suprapharyngobranchials) (Coates *et al*., 2018), *Ptomacanthus* has a chain of pharyngobranchials that sit between each arch (Dearden *et al*., 2019), and *Ozarcus* has anteriorly oriented infrapharyngobranchials and suprapharyngobranchials (Pradel *et al*., 2014). Moreover, while the pharyngobranchials are posteriorly oriented in *Acanthodes*, they lack the long swept-back blade that is present in living elasmobranchs and holocephalans (Pradel *et al*., 2014; Dearden *et al*., 2019). Despite these differences *Acanthodes*, *Gladbachus*, and *Ptomacanthus* share a chain of elements with dorsal ridges that bridge the gap between the tops of the epibranchials (Coates *et al*., 2018). *Ozarcus* is similar in the sense that the infrapharyngobranchials 3 and 4+5 have a medial ridge and a lateral shelf, while suprapharyngobranchial 1 is shaped similarly to the pharyngobranchials of *Acanthodes* (Pradel *et al*., 2014, fig. 1c). Thus these may be a common anatomy irrespective of the orientation of individual processes on the elements.

### Functional anatomy of the jaw

Due to its numerous, large pharyngeal rakers and the absence of teeth, *Acanthodes* is usually interpreted as a suspension feeder(Janvier, 1996). The orientation of the jaw articulations (Fig. 10) means that the gape would have allowed it to present a larger oral area anteriorly to take in water and food, while the numerous large gill rakers would have provided an effective separation medium (Hamann & Blanke, 2022). Increasing the gape laterally may have been particularly important due to the anguilliform body shape of *Acanthodes* giving it a comparatively small head-on profile relative to its body size. In the ecological context of the late Palaeozoic *Acanthodes* is part of a wider trend towards the evolution of eel-like body shapes in a phylogenetically diverse range of fishes (Stack *et al*., 2020).

It is becoming apparent that the mid-late Palaeozoic saw the evolution of diverse specialized feeding strategies in chondrichthyans. These include suction feeders (Coates *et al*., 2019; Dearden *et al*., 2023), grasping set-ups (Frey *et al*., 2020), and nektonic suspension feeders (Coates *et al*., 2018). Although they foreshadow some modern elasmobranch feeding strategies, these all pre-date the evolution of the specialized jaw suspensions upon which diverse modern elasmobranch feeding strategies are based (Maisey, 1980). Among these *Acanthodes* is especially noteworthy as although *Acanthodes confusus* is Permian, its skeleton is almost identical to taxa from the middle Devonian (Burrow *et al*., 2009, 2012). This includes in aspects of the specialised mandibular apparatus: the expanded anterior end of the Meckel’s cartilage appears to be present in *Halimacanthodes* (Burrow *et al*., 2012; Dearden & Giles, 2021), while a double-faceted palatoquadrate has been reported in *Cheiracanthus* (Miles, 1973a). Although *Acanthodes* is often characterised as the last gasp of the doomed acanthodians, it is perhaps better framed as the last member of a 100 million year-old lineage with a success based in an innovative feeding mechanism.

## Acknowledgements

Many thanks to Nathalie Poulet-Crovisier and Florent Goussard (MNHN CR2P) for help with 3D segmentation and visualisation. Also thanks to Marta Bellato and Patricia Wils at AST-RX, platform for Scientific Access to X-ray Tomography, UMS 2700 2AD Data Acquisition and Analysis for Natural History CNRS-MNHN, Paris.

## CRediT Statement

**RPD:** Conceptualisation, Investigation, Data Curation, Writing – Original Draft, Writing – Review & Editing, Visualisation, Funding acquisition. **AH:** Conceptualisation, Writing – Review & Editing, Funding acquisition. **AP:** Conceptualisation, Investigation, Writing – Review & Editing, Funding acquisition.

## Supplementary Data

Supplementary data comprises one supplementary table (Supplementary Table 1)

---

**Supplementary Table 1.** Comparison of terminologies for parts of the branchial skeleton of *Acanthodes confuses*.

| This study | Reis (1886) | Watson (1937) | Nelson (1967) | Miles (1973) | Jarvik (1977) | Gardiner (1984) |
| --- | --- | --- | --- | --- | --- | --- |
| Basihyal (anterior) | Linguale Copula des Hyoidbogens | basi-hyal | basibranchial 1 | basibranchial | - | basibranchial |
| Basihyal (posterior) | Copulae der Kiemenbogen | basi-branchials | basibranchials | basibranchial | - | basibranchial |
| Ceratohyal (ventral) | Praehyoid | hypo-hyal | hypohyal | ceratohyal | ceratohyal | ceratohyal |
| Ceratohyal (dorsal) | Hyoid | hypo-branchial | ceratohyal | ceratohyal | ceratohyal | ceratohyal |
| Interhyal | - | - | - | interhyal | - | - |
| Hyomandibula (ventral) | Hyomandibel | epi-hyal | epihyal | hyomandibula | hyomandibula | - |
| Hyomandibula (dorsal) | Epihyomandibulare | - | pharyngo-hyal | hyomandibula | hyomandibula | - |
| Hypobranchial | Hypobranchiale | - | ?hypobranchial | hypobranchial | - | hypobranchial |
| Ceratobranchial (ventral) | Ventrale Segmente der Kiemenbogen | hypo-hyal | hypobranchial | ceratobranchial | ceratobranchial | ceratobranchial |
| Ceratobranchial (dorsal) | Ventrale Segmente der Kiemenbogen | cerato-branchial | ceratobranchial | ceratobranchial | ceratobranchial | ceratobranchial |
| Epibranchial (ventral) | Dorsales Segmente der Kiemenbogen | epi-branchial | epibranchial | epibranchial | epibranchial | - |
| Epibranchial (dorsal) | Dorsales Segmente der Kiemenbogen | pharyngo-branchial | epibranchial | epibranchial | epibranchial | - |
| Pharyngobranchial | Pharyngobranchiale | - | pharyngobranchial | pharyngobranchial | infrapharyngobranchial | - |

